# Evolutionary study of the *412/mdg1* lineage of the *Ty3/gypsy* group of LTR retrotransposons in Diptera

**DOI:** 10.1101/2020.09.24.311225

**Authors:** Natasha Avila Bertocchi, Fabiano Pimentel Torres, Maríndia Deprá, Vera Lúcia da Silva Valente

**Author notes:** Corresponding author, (VSL).

## Abstract

LTR-retrotransposons are structurally similar to retroviruses, as they possess the enzymes reverse transcriptase, Ribonuclease H, integrase, proteinase, and the *gag* gene and are flanked by long terminal repeats (LTRs). The *412/mdg1* lineage, belonging to the *Ty3/Gypsy* group, consists of the TEs *412, mdg1, stalker, pilgrim*, and *blood*. The *412/mdg1* lineage is distinguished from the others in the *gypsy* group in that it has small ORFs at the beginning of the TE and is highly similar to the *pol* ORF among the TEs that make up the lineage. In this study, our aim was to elucidate the evolutionary history of the *412/mdg1* lineage in the 127 dipteran genomes available to date, and the characteristics of the sequences in each genome. We used the canonical TE *412* probe described in *Drosophila melanogaster* as the query. We found sequences homologous to the *412/mdg1* lineage restricted to the suborder Brachycera. These sequences are widely distributed in drosophilids but are also present in other groups of flies. We note the presence of the *412/mg1* lineage in tsetse flies (*Glossina*). Furthermore, our results showed an elaborate evolutionary history for the *pol* ORF in the *412/mdg1* lineage of the LTR-Retrotransposon.

## Introduction

Transposable elements (TEs) are part of the repetitive fraction of the genome capable of mobilizing within and between genomes. These characteristics of mobilization and parasites in genomes make TEs the most abundant and ubiquitous sequences in nature, influencing the evolutionary trajectory of host genomes in different ways [1]. TEs may be involved in diseases such as retrotransposon *L1* in the development of cancers in humans [2]; in chromosomal changes such as the *Galileo* element, which causes chromosomal inversions in *Drosophila buzzatii* [3]; or even in influencing the expression of genes, such as the *accord*-like element in the 5’ region of cytochrome P450, which confers insecticide resistance in *D. melanogaster* [4]. TEs also influence the size of the genomes and constitute the largest fraction of eukaryotic genomes. TEs may comprise 65% of the genome of insects, as in the migratory locust (*Locusta migratoria*) agricultural pest in Africa [5].

TEs can be divided according to the mechanism and intermediate transposition molecule [6–8]. Retrotransposons have the system called ‘copy-and-paste’ and RNA as an intermediate mobilization, and DNA transposons with the ‘cut-and-paste’ system and DNA as an intermediate molecule for transposition. Retrotransposons can be subdivided into two large groups: retrotransposons without long terminal repetitions (LTRs), and retrotransposons with LTRs. Both groups of retrotransposons share their evolutionary history with viruses, but only LTR-retrotransposons share an evolutionary history with retroviruses [9].

Retroviruses, endogenous retroviruses (ERVs), and retrotransposons have a common origin. All three must be transcribed back into DNA to insert/excise in the host genome, for which they use a similar enzyme structure. Retroviruses infect vertebrates and if there is integration with the host’s chromosomes in the germ cells allowing them to perpetuate in the species, they become the so-called endogenous retroviruses (ERVs) [10]. In the genomes of non-vertebrate eukaryotic organisms, retrovirus-like are called LTR-retrotransposons [9].

LTR retrotransposons have similar characteristics to retroviruses, such as LTRs flanking the open reading frames (ORFs) that code for proteinase (PR), reverse transcriptase (RT), Ribonuclease H (RH), integrase (INT), and *gag*-like proteins; and are organized similarly to retroviruses [11,12]. However, only a few LTR retrotransposons have putative *env* genes, responsible for the viral envelope [9]. The *gag* gene (ORF1) is responsible for the virus-like particle (VLP), and the *pol* gene (ORF2) is responsible for and the other enzymes (PR-RT-RH-INT) that are responsible for the mobilization. LTR-retrotransposons are mobilized via a mechanism that requires reverse transcription of mRNA. The 5’ LTR has a promoter that is recognized by the host’s RNA pol II, after transcription of the intermediate mRNA, which is exported to the cytoplasm where the translation occurs. The VLP encapsulates the mRNA, which is reverse-transcribed, giving rise to the cDNA. The latter is then sent to the nucleus, where it will be inserted into the host genome by the action of integrase [13].

The phylogenetic relationships of LTR-retrotransposons subdivide them into two groups, called *Ty1/copia* and *Ty3/Gypsy* [14]. The phylogenetic tree of the *Ty3/Gypsy* group has two large branches, based on the *pol* ORF (PR-RT-RH-INT) [15]. The first branch, called chromoviruses because they contain a chromodomain in the integrase, consists of TEs of plants, fungi, and vertebrate animals [14–16]. The second branch, called non-chromoviral, consists of TEs of insects and other groups of organisms [14,15,17,18]. Inside the *Ty3/Gypsy* non-chromoviral branch is the *412/mdg1* lineage [17].

The *412/ mdg1* lineage consists of the *412, stalker, pilgrim, blood*, and *mdg1* elements, characterized in the genome of *D. melanogaster* [17]. Retrotransposon *412* was the first of the *412/mdg1* lineage to be described, with approximately 7000 base pairs (bp) [19,20]. Retrotransposons of the *412/ mdg1* lineage share structural features such as two small ORFs (sORFs) within the 5’ region, ORF1 (*gag*), and ORF2 (*pol*) [14,17,19]. Putative sORFs are a distinctive feature of the *412/mdg1* lineage, but still have no known role in the survival of the element [21,22]. However, copies of *blood* without sORFs showed greater competitive potential compared to copies with sORFs in natural populations of *D. melanogaster* [22]. In addition, when analyzed separately, the sORFs, ORF1 and ORF2, showed differences in phylogenetic relationships between the elements of the lineage [17]. However, the same phylogenetic relationships among the elements of the lineage were observed in the alignment of each of the different domains (PR-RT-RH-INT) of ORF2 [17].

Historically, the reconstruction of the phylogenetic relationships of LTR-retrotransposons is based on the RT domain, as it is one of the largest and most conserved domains. These evolutionary relationships were later supported with the addition of the other ORF2 domains [9,15,23–26]. In addition, the RT and RH domains have been shown to provide a good resolution of the phylogenetic relationships within the *Ty3/gypsy* group [24].

Within Insecta, the order Diptera is the best characterized in relation to the fraction of transposable elements (TEs); however, most studies have examined model organisms such as species of the genus *Drosophila* or mosquito disease vectors such as *Aedes aegypti* [27,28]. The percentage of TEs varies widely among dipteran genera, ranging from 1% in the fly *Belgica antarctica* (restricted to Antarctica) to more than 50% in *Aedes aegypti* [27,29]. Within the TEs, the order of retrotransposons constitutes the largest fraction of TEs in dipteran genera [27,28,30]. Our research group has been studying the presence of TEs, including *412* in Neotropical species of *Drosophila*, where we supposed *a priori* that the wide diversity of biomes and environments may have provided many opportunities for evolutionary adjustments in these genomes. Blauth et al. [31], comparatively analyzing *412* of *D. melanogaster* and *D. willistoni*, observed that *D. willistoni* showed more restricted hybridization patterns during embryogenesis, probably due to the difference in sequence similarity between the canonical TE of *D. melanogaster* and the TE in the *D. willistoni* genome. The present study aimed to extend the investigation of the TEs of Neotropical species and/or other native or introduced species of flies, exploiting newly available methods.

The increase in genome sequencing provides an extraordinary opportunity to deepen knowledge about the evolution of TE lineages such as *412/mdg1*, and to fill the gaps in the distribution and phylogeny of their main lineages in the dipteran genomes. In this study, we peovie evidenve the conservation of the enzymes that constitute ORF2 in the *412/mdg1* lineage in the genomes of different evolutionary lines of flies. We have possibly identified a new branch in the *412/mdg1* lineages in the species of the genus *Glossina* (tsetse), the vector of the parasite that causes human sleeping sickness. Furthermore, we elucidated the intricate nature of the evolutionary relationships involving the ORF2 sequences of the *412/mdg1* lineage.

## Material and Methods

### Identification of *412* sequences from dipteran genomes

The 127 publicly available dipteran WGS (whole-genome sequences) from the National Center for Biotechnology Information (NCBI) and the *Drosophila buzzatii* Genome Project (https://dbuz.uab.cat/welcome.php) were used in this study, and were last accessed in June 2019. The query utilized was *412* (GenBank accession number X04132) sequence from *D. melanogaster* available from NCBI.

*412* query was used for BLASTn searches in the dipteran species WGS in the NCBI database [32]. A list of the species analyzed and the corresponding sequence data is provided in Table 1. The sequences with an E-value lower than e-10 were extracted. In Table 1: column “BLAST_*412* query” are the genomes with positive results in BLASTn, and column “*412*-like” are the genomes with homologous sequences after refinement (described below).

**Table 1:**
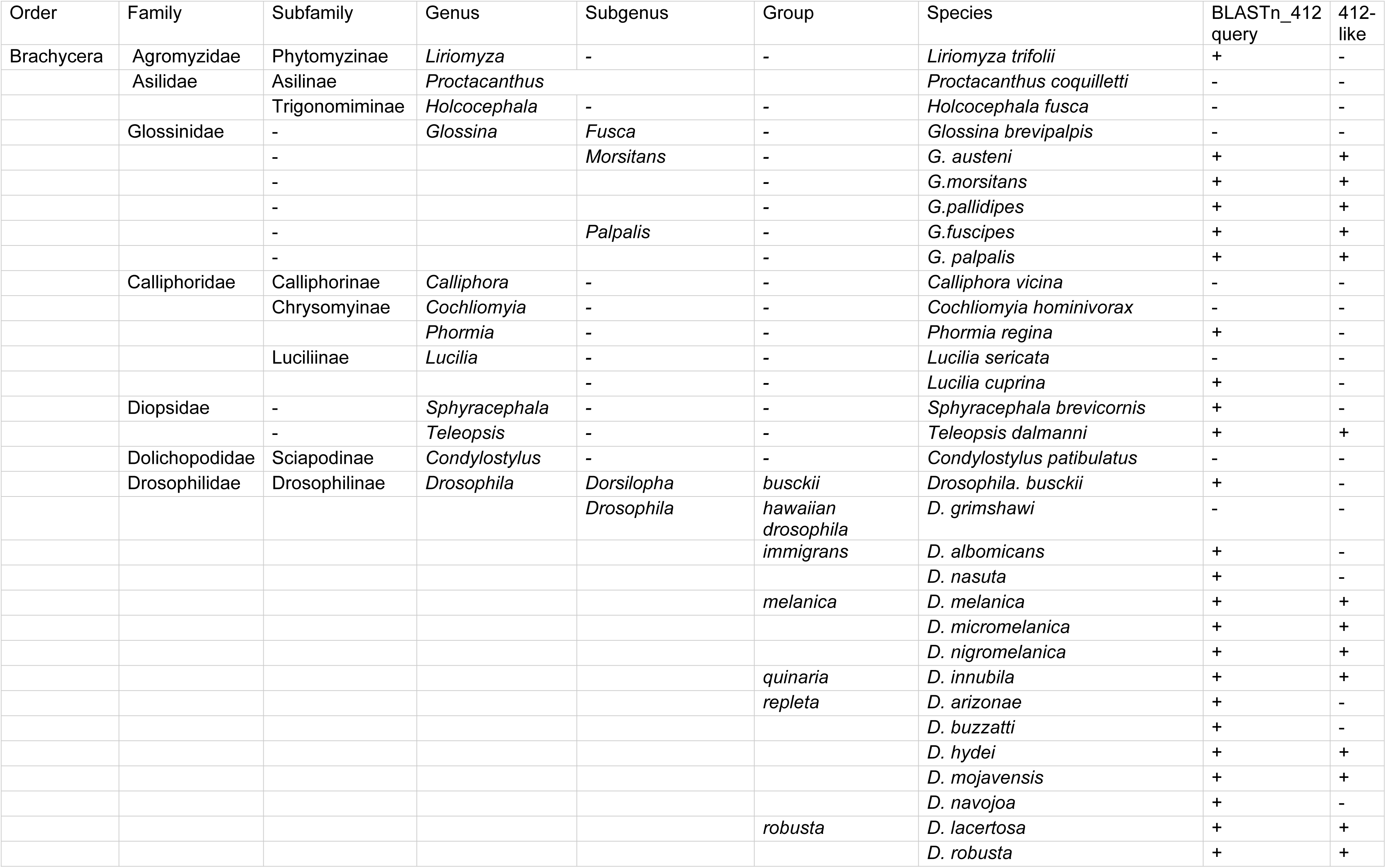
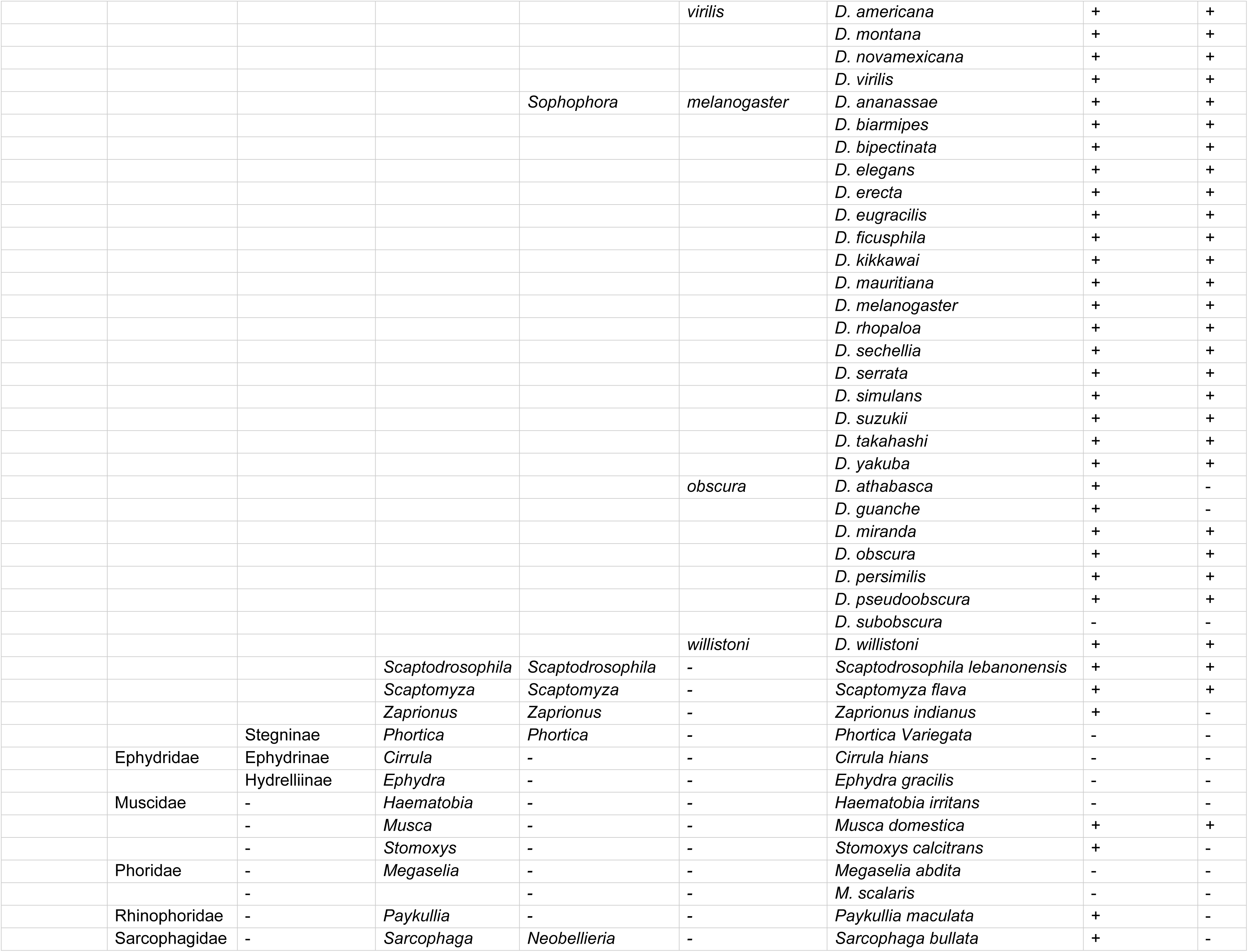
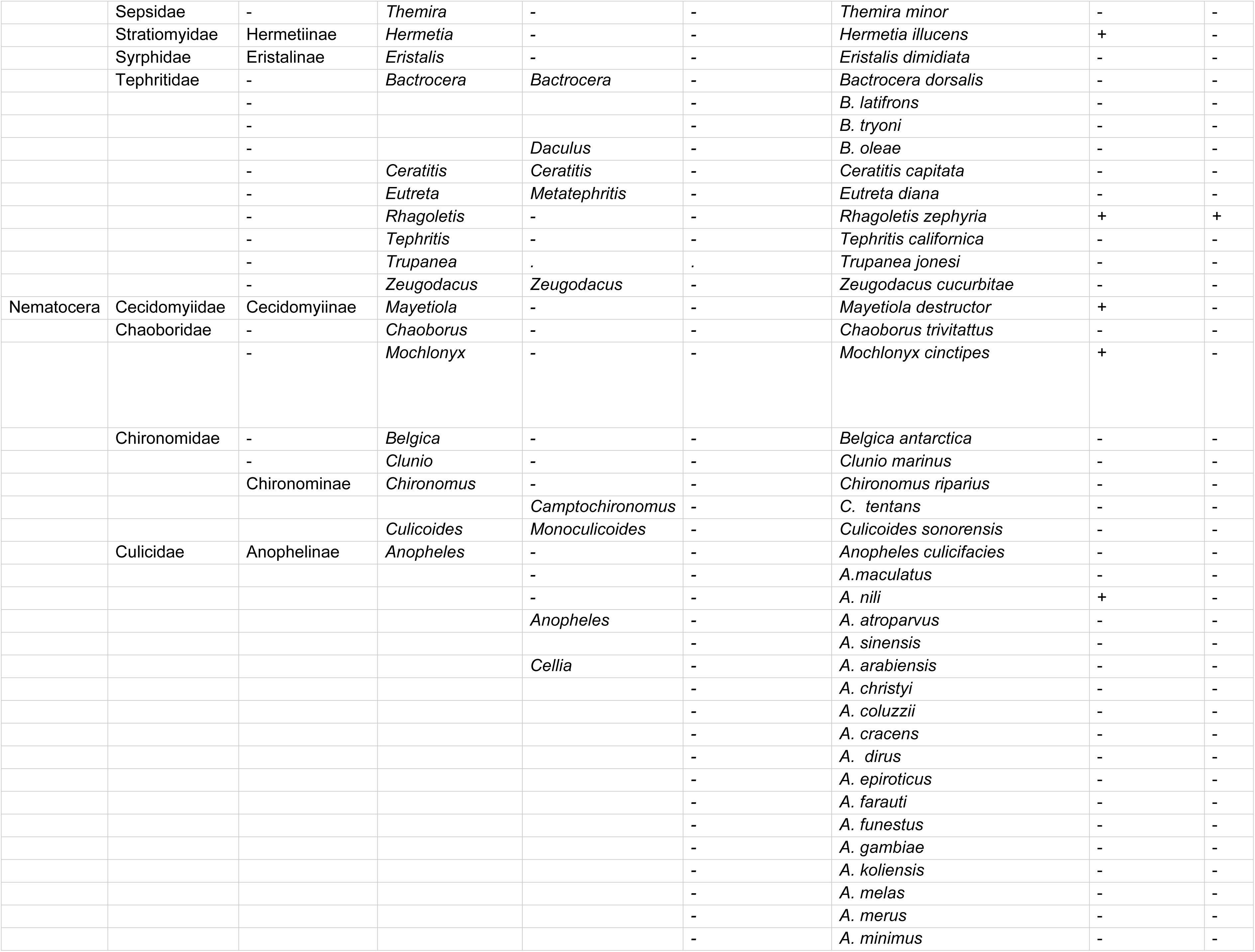
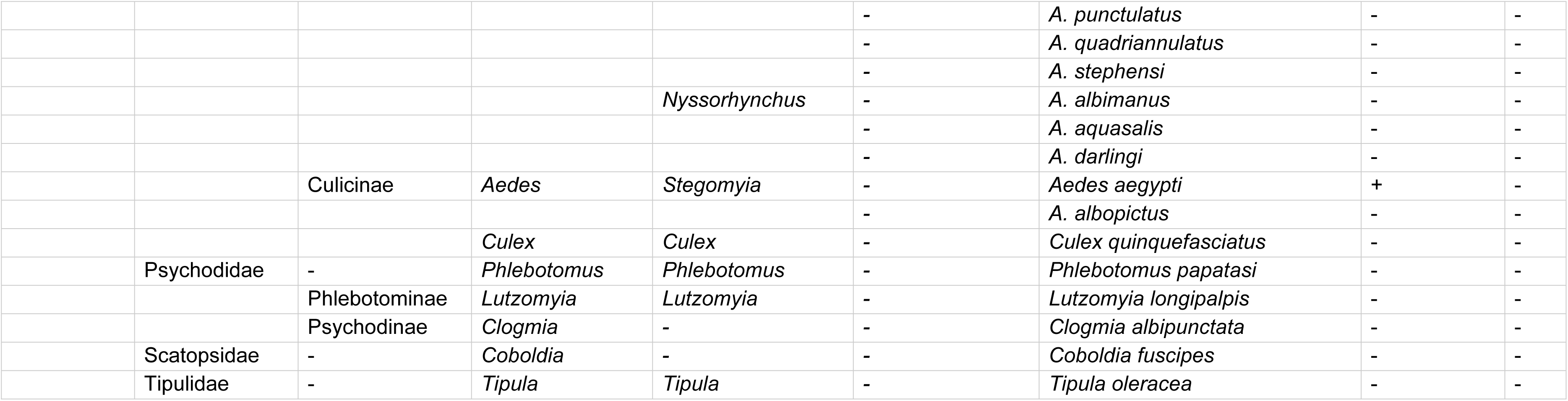
Presence/absence of *412*-like sequences in the genomes of Order Diptera. (+) *412*-like elements present; (-) *412*-like elements absent.

### Sequence analysis

The genome definition with the presence of sequences homologous to *412* was carried out with two stages of refinement. In the first stage, the homologous sequences in each genome were aligned using the MAFFT software with the default parameters, adjusting the direction according to the complete TE *412* reference sequence [33]. In this first stage, sequences that covered most of the element (*412* query) or different segments of the TE were retained, making it possible to establish a consensus sequence for each genome. The consensus sequences were obtained using the UGENE software, with the default parameters showing the most common nucleotide in each position [34]. In the second stage, the previously generated consensus sequences were aligned again by the MAFFT software, using the G-INS-1 method and the direction of the reference sequence [33]. The alignments were refined manually when necessary and the sequences were translated using the Aliview software. Genomes containing *412*-like elements were considered to be those that contained sequences with at least 50% similarity at the level of nucleotides and amino acids with the sequence of the internal region of the *412* element (X04132) (S1 Table). These are sORFs, the *gag* gene (ORF1), and the *pol* gene (ORF2), as originally described [19].

### Phylogenetic analyses

Conserved ORF2 domains (RT-RH) were used in the phylogenetic analyses as described by [25]. The domains were identified using the Conserved Domain Database (CDD) platform https://www.ncbi.nlm.nih.gov/Structure/cdd/wrpsb.cgi. To confirm the relationship of the sequences obtained with the elements of the *412/mdg1* lineage, the following *D. melanogaster* sequences were included in the analysis: *412* (GenBank accession number X04132), *mdg1* (GenBank accession number X59545.1), *pilgrim* (GenBank accession number AC007146), and STALKER4_I and BLOOD_I from the Repbase database. Element *17.6* (GenBank number X01472.1) was used as the outgroup. Bayesian phylogenetic analysis was performed with BEAST v1.10.4 software, using the LG model [35] with gamma distribution and invariant sites, through Cipres Computational Resources [36]. The Markov Chain Monte Carlo (MCMC) was run for 10,000,000 iterations, with trees saved every 10,000 iterations, after a 2,500 burn-in.

The phylogenetic reconstruction of the analyzed species was based on the nuclear gene *Amyrel*. Sequences were downloaded using the access number provided by Tambones et al. [37] (S2 Table). Additional sequences were obtained directly from NCBI, using *D. melanogaster* as a query or another related species when no results were recovered. Bayesian inference was performed with BEAST v1.10.4 software, using General Time Reversible (GTR) with gamma distribution and invariant sites, through Cipres Computational Resources [36]. The mosquito species *Anopheles gambiae* was used as the outgroup.

### Median-joining networks

Diagrams using Median-joining were obtained from conserved regions used in previous phylogenetic reconstructions (conserved domains RT-RH). The alignments were converted to the Roehl format, with deletion of the invariant sites using the DNASP version 6 program, and the diagrams were generated using the Fluxus Network software, with default parameters [38,39].

## Results

### Genomes with *412-*like retrotransposon

The presence of *412*-like sequences was evaluated in 127 available genomes of the order Diptera: 87 genomes of suborder Brachycera (flies) and 40 genomes of suborder Nematocera (mosquitoes) (Table 1). Initially, the preliminary results identified *412*-like sequences in 65 genomes, 61 in Brachycera and only 4 in Nematocera (Table 1). However, the sequences from species of Nematocera and some species of Brachycera, after refinement, proved to be quite divergent from the *412*-like sequences and were excluded (Table 1 and S1 Table).

Within the suborder Brachycera, sequences were identified in the families Tephritidae (1 species), Glossinidae (5 species), Muscidae (1 species), Diopsidae (1 species) and Drosophilidae (36 species). Thus, the *412*-like sequences showed a wide distribution in the subfamily Drosophilinae, including the genera *Scaptodrosophila*, *Scaptomyza*, and mainly *Drosophila* (Table 1).

### Sequences homologous to lineage *412/mdg1*

The complete sequence of the internal region (sORFs, ORF1, and ORF2) of *412*-like were detected only in the *Drosophila* genome, in *D. melanogaster*, *D. mauritiana*, and *D. simulans* (Fig 1). In *D. sechellia* a complete sequence of *412* was not identified and all copies had some deletions; however, it was possible to obtain a consensus sequence from an assembly of these sequences (Fig 1). The copies detected in these genomes were identical or nearly so to the reference *412* (accession number X04132) at the nucleotide level, ranging from 94% in *D. sechellia* to 100% in *D. melanogaster* (S3 Table).

**Figure 1:**
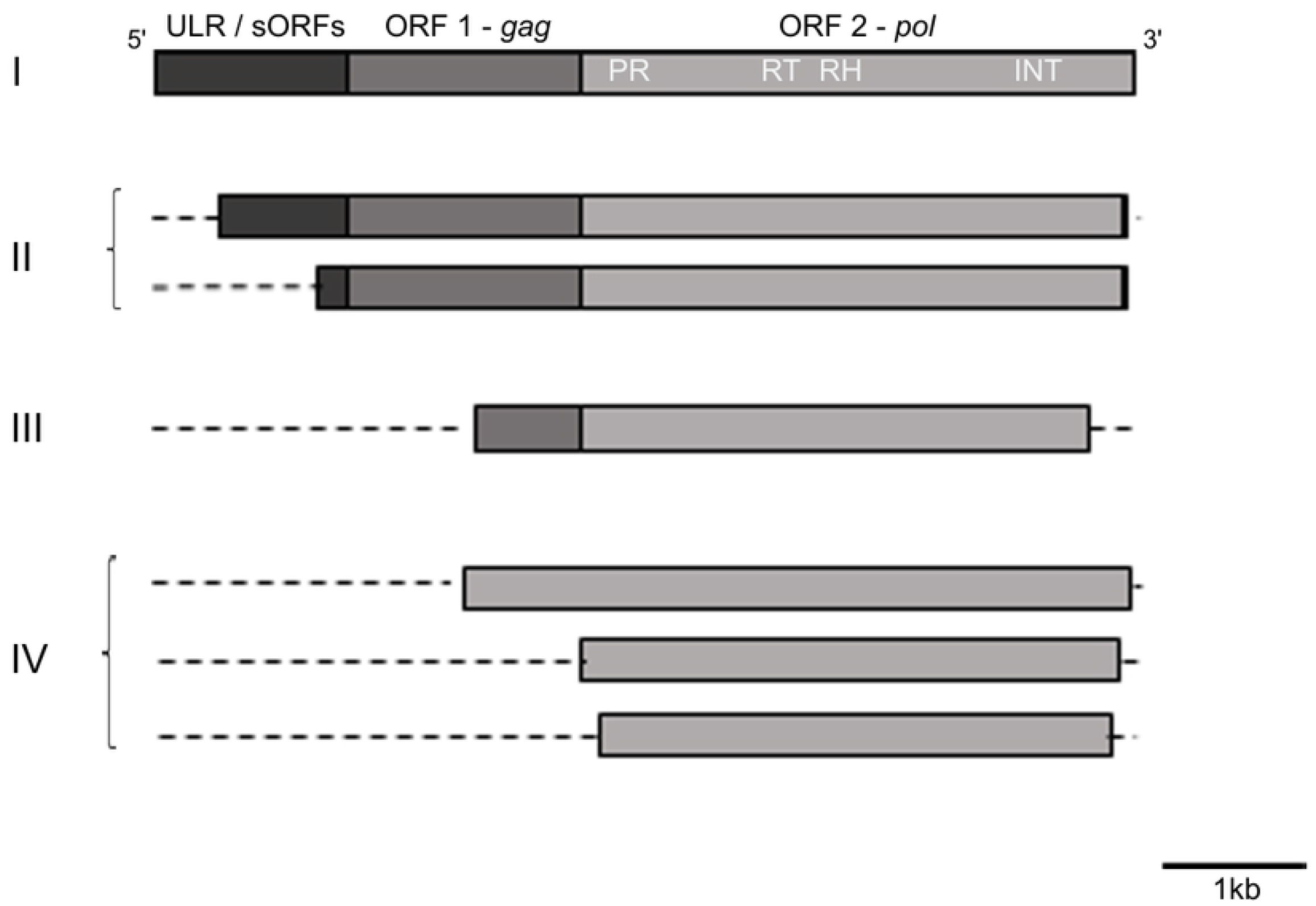
Schematic representation of the reconstructed *412-like* copies compared to the *412* canonical. The gray rectangles indicate an untranslated leader region (ULR) and two small putative ORFs (sORFs), the *gag* gene (ORF1) and the *pol* gene (ORF2). Different domains of the ORF2 are indicated as follows: protease, PR; reverse transcriptase, RT; Ribonuclease H, RH; integrase, INT. Species were divided into four groups according to their structural characteristics. Groups I and II are Family Drosophilidae, subgenus *Sophophora* – *melanogaster* group. **I** – *412* (X04132), *Drosophila melanogaster, D. mauritiana, D. sechellia, D. simulans*. **II** – *D. erecta* and *D. yakuba*. **III** – Subgenus *Drosophila* – *virilis* group: *D. virilis;* **IV** – Family Glossinidae, subgenus *Morsitans*: *Glossina austeni, G. morsitans, G. pallidipes;* Subgenus *Palpalis*: *G. fuscipes, G. palpalis;* Family Diopsidae: *Teleopsis dalmanni;* Family Drosophilidae, subgenus *Drosophila – melanica* group*: D. melanica, D. micromelanica, D. nigromelanica, D. innubila; repleta* group: *D. hydei, D. mojavensis; robusta* group*: D. lacertosa, D. robusta; virilis* group: *D. americana, D. montana, D. novamexicana;* Subgenus *Sophophora – melanogaster* group: *D. ananassae, D. biarmipes, D. bipectinata, D. elegans, D. eugracilis, D. ficusphila, D. kikkawai, D. rhopaloa, D. serrata, D. suzukii, D. takahashi; obscura* group: *D. miranda, D. obscura, D. persimilis, D. pseudoobscura; willistoni* group: *D. willistoni;* Subgenus *Scaptodrosophila*: *Scaptodrosophila lebanonensis;* Subgenus *Scaptomyza*: *Scaptomyza flava;* Family Muscidae: *Musca domestica;* Family Tephritidae: *Rhagoletis zephyria*

Other regions of the TE in the remaining genomes, analyzed only in the species *D. erecta*, *D. yakuba*, *D. virilis*, *D. suzukii*, and *Scaptomyza lebanonensis*, possess sequences partially homologous to ORF1 of *412*. In addition, all genomes analyzed showed sequences homologous to ORF2 of *412*. The similarity of the sequences in relation to the query, level of amino acids, ranged between 100% in *D. melanogaster* and 53% in *D. nigromelanica* (Fig 1 and S1 Table). Among the enzyme genes that constitute ORF2, the most conserved sequences were related to RT and RH.

### Evolutionary analysis of the sequences evaluated

The searches for the dispersion of *412* in the dipteran genomes showed that the elements of the *412/mdg1* lineage are very similar to each other, as seen in Figures 2 and 3. We reconstructed the phylogenetic relationships of the *412/mdg1* lineages based on a fragment of ORF2, the conserved regions of the RT, and RH genes from the consensus sequences obtained. The consensus sequences represent an approximation of the genes that gave rise to the visible copies in the genomes. The resulting tree showed a topology with the species of flies, with emphasis on *Drosophila*, not grouping according to its host genome, as observed in the phylogeny of the *Amyrel* gene (S1 Fig). *Musca domestica*, *R. zephyria*, and *T. dalmani* were positioned in more external branches (Fig 2). Lineages *412*, *stalker*, and *pilgrim* formed three distinct arms, where most species were grouped (Fig 2). However, the lineages *mdg1* and *blood* were placed in another outer arm, distant from the other lineages and shared with the *Glossina* sequence.

**Figure 2:**
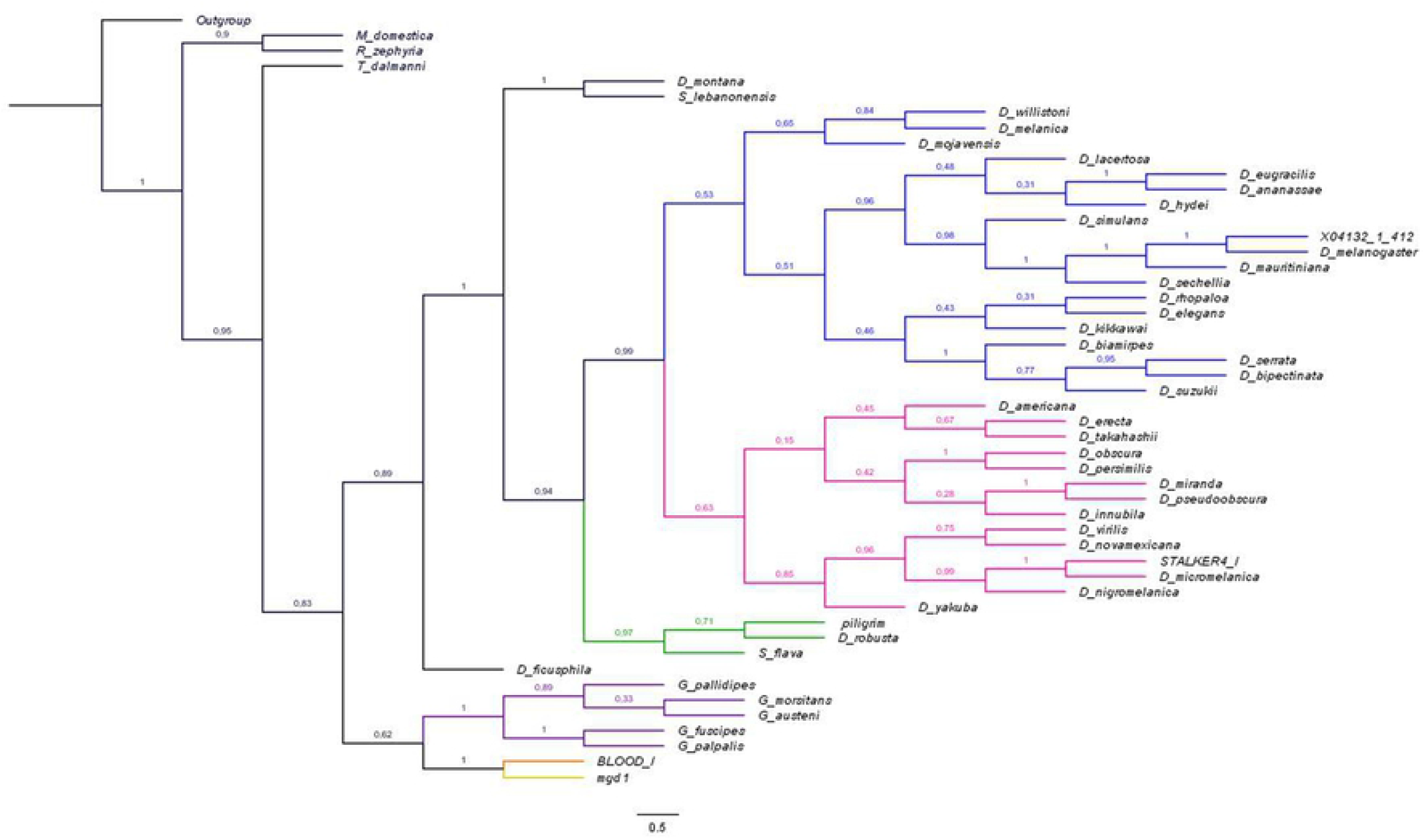
Phylogenetic reconstructions of the *412/mdg1* lineage based on reverse transcriptase (RT) and Ribonuclease H (RH) regions. Bayesian phylogenetic analysis was performed with BEAST v1.10.4 software [Cipres Computational Resources [36]], using the LG+G+I model. The Markov Chain Monte Carlo (MCMC) was run for 10,000,000 iterations, with trees saved every 10,000 iterations, after a 2,500 burn-in. Lineage *412/mdg1: 412* (blue), *stalker* (pink), *mdg1* (yellow), *pilgrim* (green), *blood* (orange), *Glossina* species (purple), and median vectors (red). Element 17.6 was used as the outgroup.

**Figure 3:**
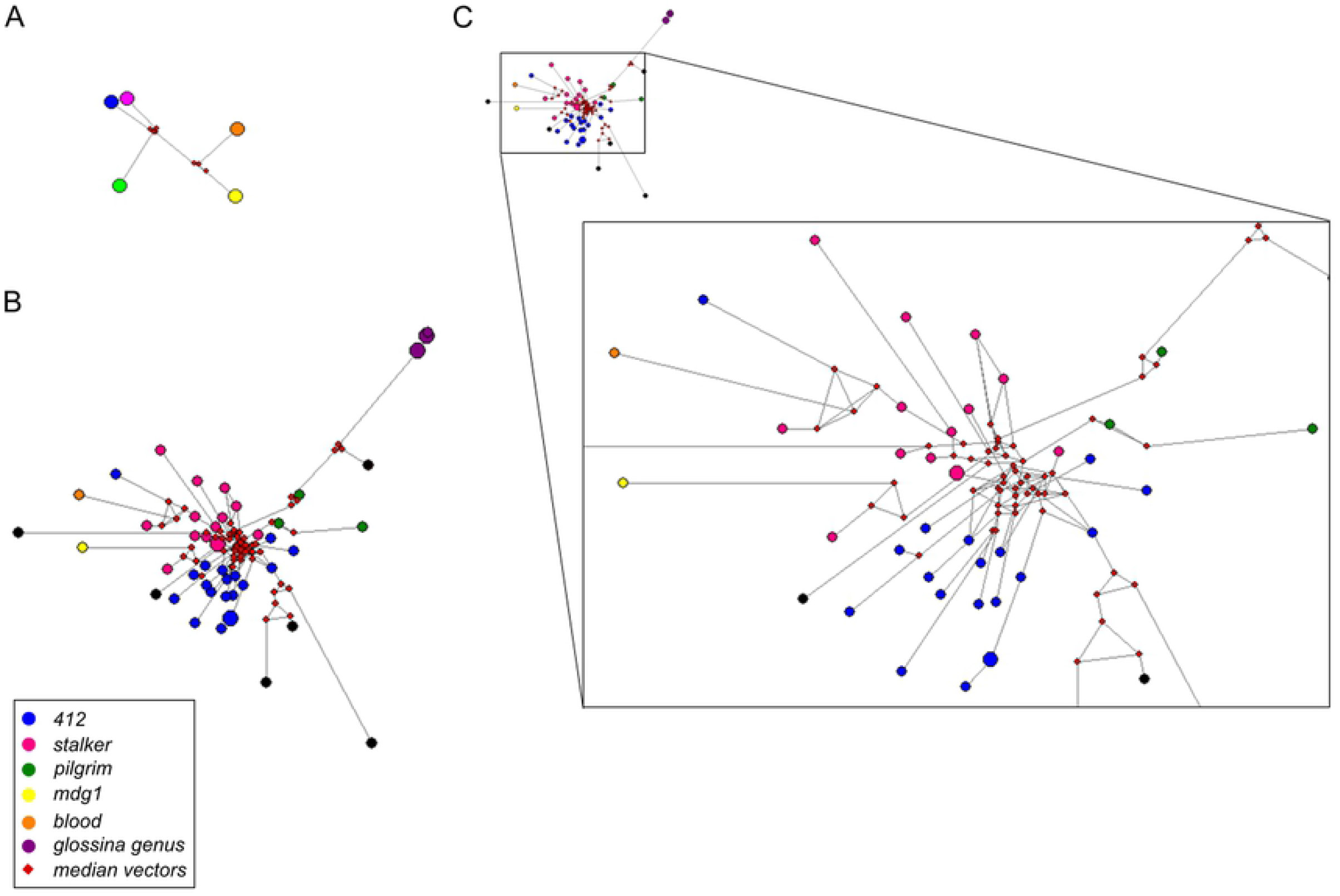
Network reconstructions. Median-joining network analyses for the *412/mdg1* lineage. The size of each circle indicates the number of sequences grouped together. (a) Network for the elements in the *412/mdg1* lineage. Each element in the lineage is differently colored. (b) The network of all sequences used in the phylogenetic reconstructions. (c) Detail of the *412/mdg1* lineage relationships. The sequences are colored according to the clustering in the phylogenetic tree.

We built a network with the same consensus sequences used in the phylogenetic tree, as a new approach to analyzing the complex relationship between the different elements that comprised the *mdg1/412* lineage and consequently shaped its evolution in different genomes. Inspection of the network shows that the elements *412*, *stalker*, and *pilgrim* are more closely related to each other than to *mdg1* and *blood* (Fig 3A). Analysis of all the sequences reveals the intricate relationship among the sequences, demonstrated by the central median vectors, which form an intricate network that links all the sequences (Fig 3B-C). On the other hand, *412* and *stalker* are not closely related to the phylogenetic tree but are inside sister groups.

The different approaches used to unveil the evolution of this group of TEs, phylogenetic tree and network, indicated that the sequences obtained in the five genomes that showed positive results in the BLAST of members of the genus *Glossina* formed a new group within the lineage *412/mdg1* (Figs 2 and 3).

## Discussion

Our results showed that the sequences homologous to *412* in the order Diptera were restricted to the suborder Brachycera, which is composed of flies (Table 1). Dipterans diverged more than 350 million years ago (mya), and at approximately 300 mya divided between flies (Brachycera) and mosquitoes (Nematocera) [40]. Therefore, we have three possibilities for the elements of the *412/mdg1* lineage: 1) they remained in the fly branch; 2) they were lost in the mosquito group; 3) or they were inserted into the fly genome after the group diverged at approximately 300 mya.

In suborder Brachycera, the *412*-like elements are widely distributed in the available genomes (Fig 1). However, complete sequences of *412* were available only for *Drosophila melanogaster, D. mauritiana, D. simulans* and *D. sechellia* (consensus) (Fig 1). We found complete sequences of *412*, including integral LTRs (data not shown), only in closely evolutionarily related *Drosophila* species. *D. melanogaster* appeared between 2.5 and 3.4 Mya, and *D. simulans, D. sechellia*, and *D. mauritiana* diverged at approximately 0.5 Mya, forming the *melanogaster* supercomplex [41]. Furthermore, we also detected partially complete *412* copies in the genomes of *D. yakuba, D. erecta*, and *D. ananasse*, in the first two copies was missing the region of the sORFs, and in the last one, fragments of ORF2 were detected (Fig 1). Dias et al. [42] also identified complete sequences of *412* in these same species, except for *D. mauritiana* (did not analyze), and in addition to these were also identified in *D. yakuba, D. erecta* and *D. ananasse*.

In the phylogenetic tree, the members of the families Muscidae, Tephritidae, Diopsidae, Drosophilidae, and Glossinidae are divided into different branches (Fig 2). However, inspection of the internal branches shows that elements *412, stalker*, and *pilgrim* form groups with different species regardless of the host genome (Figs. 2 and 4). The discrepancies between host genomes and the elements are more evident in the elaborate network formed by the *412/mdg1* lineage (Fig 3 A-C). Costas et al. [17], characterizing the *412/mdg1* lineage, observed differences in the phylogenetic relationships between the elements when the ORF1 and ORF2 were analyzed separately; these differences can be explained by the independent evolution of each of the conserved domains of the TEs [43]. However, did not identify differences in the relationships of the elements when analyzing the ORF2 domains individually [17].

The intricate relationships between the different elements of the *412/mdg1* lineage, with the central median vectors forming an elaborate network (Fig 3C), probably occurred due to mosaicisms of these retrotransposons. In the *412/mdg1* lineage, mosaicisms were identified within the sORFs and ORF1 between *412* and *stalker, 412*, and *mdg1* in *D. melanogaster, D. simulans* and *D. yakuba* populations [8,21]. According to Costas et al. [17], the *stalker* element is the result of recombination events in the ancestral family of what are currently the other elements of the lineage. LTR retrotransposons may be prone to genetic rearrangements (mosaicisms), as two viral genomes packaged in the same virus-like particle can recombine during the reverse-transcription step between RNA genomes [44].

Another interesting feature of our results was the homologous sequences of the *412/mdg1* lineage in *Glossina*. Of the six species analyzed, *G. palpalis, G. fuscipes, G. austeni, G. morsitans, G. pallidipes*, and *G. brevipalpis*, we not found homologous sequences of the *412/mdg1* lineage in *G. brevipalpis* (Table 1). Knowledge of the genomic characteristics and TEs in tsetse flies is still sparse and preliminary. However, the authors who sequenced and analyzed the genomes of these 6 tsetse species observed that *G. brevipalpis* is the most basal species in the group, with the fewest TEs (~ 35% of the total genome), practically without LTR-TEs, and large expansion of *mariner*-like elements [30]. Possibly the *412/mdg1* lineage was not maintained in *G. brevipalpis* as in the other species of *Glossina*. In addition, the sequences identified in the *Glossina* genomes may be another element of the *412/mdg1* lineage, possibly diversified from the *blood* and *mdg1* sequences (Figs 2 and 3). Furthermore, we identified sequences homologous to lineage *412/mdg1* in *D. persimilis* (Figs 1 and 2), which had not previously been identified [23]. Bargues and Lerat [23] initially searched the LTRs of the elements and later analyzed ORF2. In our analyses, we searched the genomes of Diptera for the internal regions of the element (sORFs, ORF1, ORF2). We identified relics of the *412/mdg1* lineage in the *D. persimilis* genome (Figs 1 and 2), but this allowed us to group this species together with other drosophilas, specifically with a sister group of *D. obscura* (Fig 2).

The *412/mdg1* lineage is widely dispersed in the genomes of flies, not only in the family Drosophilidae, and at different levels of conservation in the *pol* gene. Our search for *412* in the genomes of Diptera revealed an intricate relationship in the evolution of ORF2 between the elements *412*, *stalker, pilgrim*, *blood*, and *mdg1*. This indicates the importance of mosaicism in the evolutionary history of the *412/mdg1* lineage.

## Acknowledgements

We are grateful to M.Sc. Thays Duarte de Oliveira and M.Sc. Henrique Moreira for their valuable assistance with the figures and phylogenetic analyses.

## Supporting information

**S1 Fig: Phylogenetic relationships between sequences in the nuclear gene *Amyrel* in 44 species of suborder Brachycera.** Bayesian inference was performed with BEAST v1.10.4 software, using GTR+G+I as the substitution model, through Cipres Computational Resources [36]. Outgroup: *Anopheles gambiae*.

**S1 Table: Distance matrix of 412-like sequences**. Nucleotide and amino acid distance matrix of sequences 412-like, using UGENE software [34].

**S2 Table: Acess number *Amyrel* gene.**

**S3 Table: Distance matrix of complete copies of *412* in *Drosophila*.**

Distance matrix based on the nucleotide sequences of complete copies of *412* in *Drosophila*, using the UGENE software [34].

